# Identification of potential candidates for drug designing against *Klebsiella Pneumoniae* among hypothetical proteins of DNA Adenine Methyltransferase through subtractive genomics approach

**DOI:** 10.1101/2020.02.13.947101

**Authors:** Umairah Natasya Mohd Omeershffudin, Suresh Kumar

## Abstract

*Klebsiella Pneumoniae* is a gram-negative bacterium that is known for causing infection in nosocomial settings. As reported by WHO, this bacterial pathogen is classified as an urgent threat our most concern is that these bacterial pathogens acquired genetic traits that make them resistant towards antibiotics. The last class of antibiotics; carbapenems are not able to combat these bacterial pathogens allowing them to clonally expand their antibiotic-resistant strain. Most antibiotics target the essential pathways of the bacteria cell however these targets are no longer susceptible to the antibiotic. Hence in our study, we focus on *Klebsiella Pneumoniae* bacterial strains that contain DNA Adenine Methyltransferase domain which suggests a new potential site for a drug target. DNA methylation is seen to regulate the attenuation of bacterial virulence. In this study, all hypothetical proteins of *Klebsiella Pneumoniae* containing N6 DNA Adenine Methyltransferase domain were analysed for a potential drug target. About 32 hypothetical proteins were retrieved from Uniprot. 19 proteins were selected through a step-wise subtractive genomics approach like a selection of non-homologus proteins against the human host, selection of bacterial proteins contains an essential gene, broad-spectrum analysis, druggability analysis, Non-homology analysis against gut microbiota. Through drug target prioritization like sub-cellular analysis, drug property analysis, anti-target non-homology analysis, virulence factor analysis and protein-protein interaction analysis one drug target protein (Uniprot ID: A0A2U0NNR3) was prioritized. Identified drug target docked with potential inhibitors like are mahanine (PubChem ID: 375151), curcumin (PubChem ID: 969516), EGCG (PubChem ID: 65064), nanaomycin A (PubChem ID: 40586), parthenolide (PubChem ID: 7251185), quercetin (PubChem ID: 5280343) and trimethylaurintricarboxylic acid. Based on the moelcular docking analysis, mahanine has the highest binding affinity. In order to identify novel natural inhibitor based on mahanine fingerprint search is performed against NPASS (Natural Product Activity and Species Source databases) and Koenimbine was identified as a novel natural inhibitor based on virtual screening.

## 1. Introduction

*Klebsiella Pneumoniae* belongs to the genus of *Enterobacteriae* and classified as one of the Carbapenems-Resistant-Enterobacteriaceae (CRE) organisms. This organism causes infection in the nosocomial setting leading to a global threat as the ability of the bacterial pathogens to acquire mobile genetic traits making them resistant towards antibiotics. *Klebsiella Pneumoniae* causes a wide range of infection which includes the urinary tract infection, pneumonia and liver abscess (Paczosa 2016). The current drug targets the cellular process of the bacterial pathogens of the translation, transcription, and replication. However, the bacterial pathogens are still able to develop resistant towards the antibiotics (Petchiappan and Chatterji 2017).

The concerning emergence of this bacterial pathogen MDR has become a global threat and this MDR and as mentioned by WHO (2012). These bacterial pathogens clonally expand acquiring genetical traits allowing them to develop resistant causing an increase prevalence affecting the human population by suppressing the mortality and morbidity rate. The current antibiotics are not able to combat this bacterial pathogen allowing them to acquire mobile genetic traits that make them resistant towards the antibiotics. Although, the carbapenems and colistin are identified as the 2 most potent antibiotics however this bacteria producing resistant towards these antibiotics and identified as *Klebsiella Pneumoniae* Carbapenems(KPCs) producing (Giordano et al. 2017).

DNA Adenine Methyltransferase(Dam) is seen as a promising drug target as it involves in the epigenetics regulatory machinery that helps in sustaining the viability of the bacterial pathogens and regulates the bacterial pathogenicity (Natasya, Ismail, and Kumar 2019). Evidently, Dam is seen to regulate the modulation of that bacterial pathogen making them a promising drug target for future drug discovery (Giacomodonato et al. 2014). Hence, this study aims to identify novel antibiotic drug targets by targeting the Dam for *Klebsiella Pneumonia*.

## 2.0 Materials and Methods

This study involves several *in silico* methods that include subtractive genomic analysis, molecular docking and ADME test. The detailed workflow is summarized in Figure 1, Figure 2 and Figure 3.

**Figure 1:**
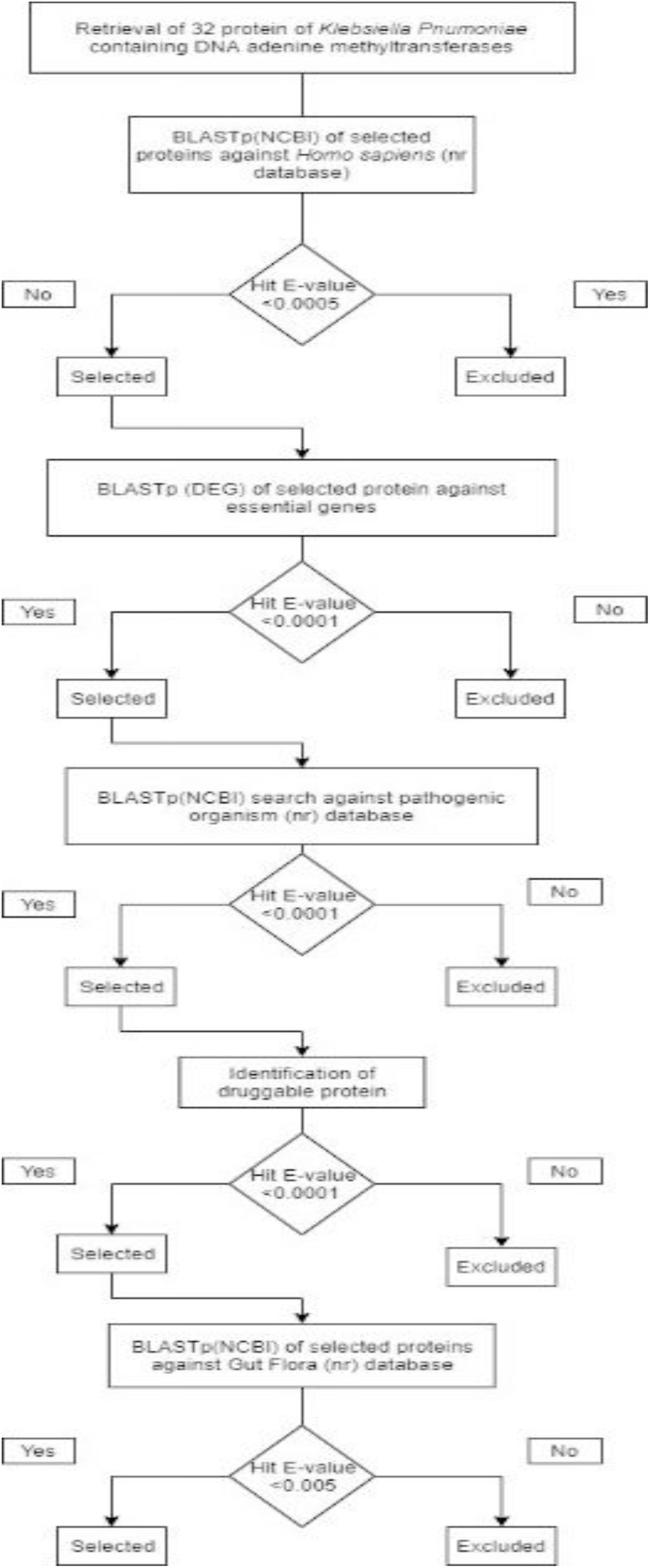
Overall workflow of identification of protein putative drug target

**Figure 2:**
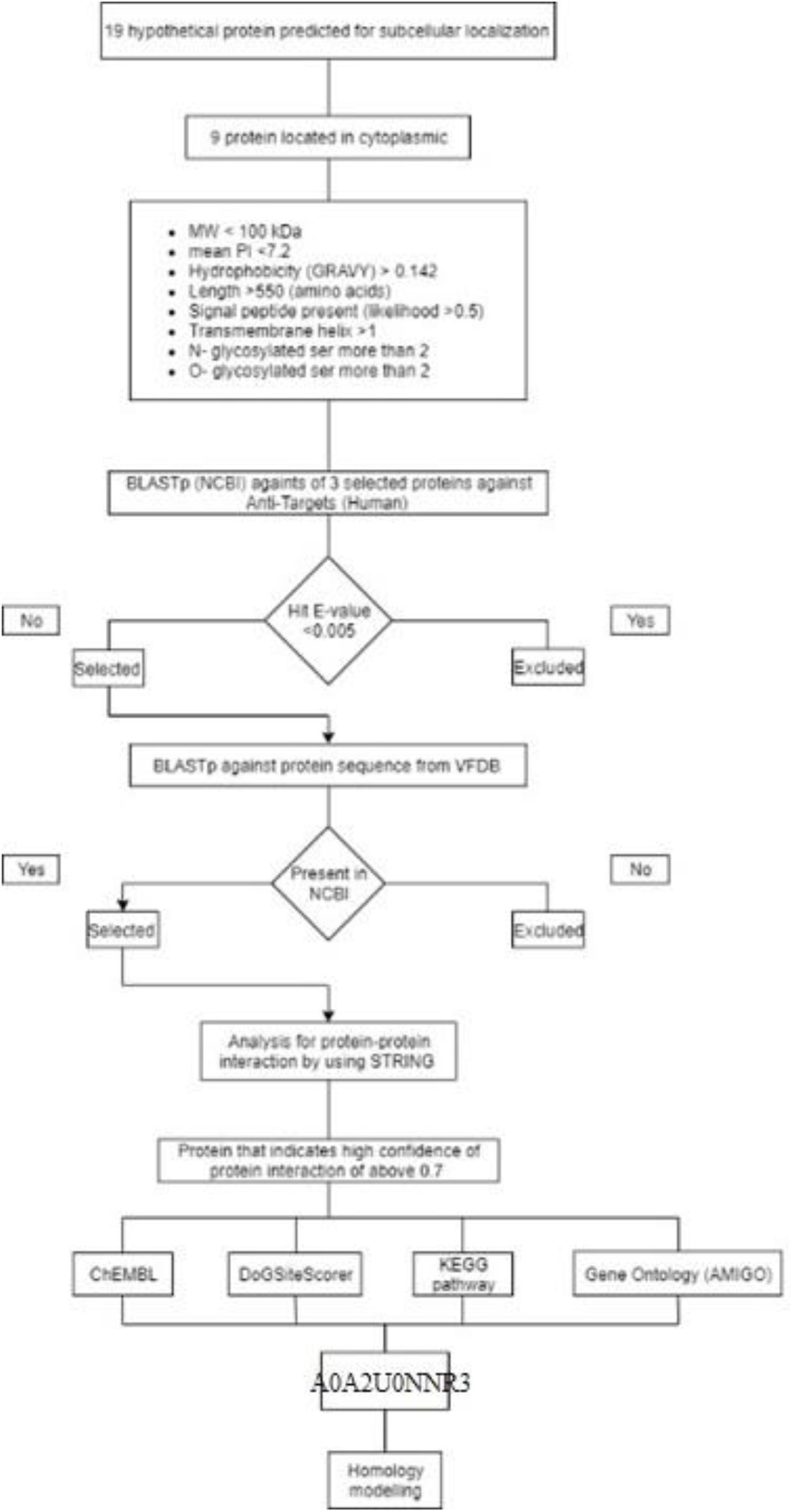
Extended workflow summarization

**Figure 3:**
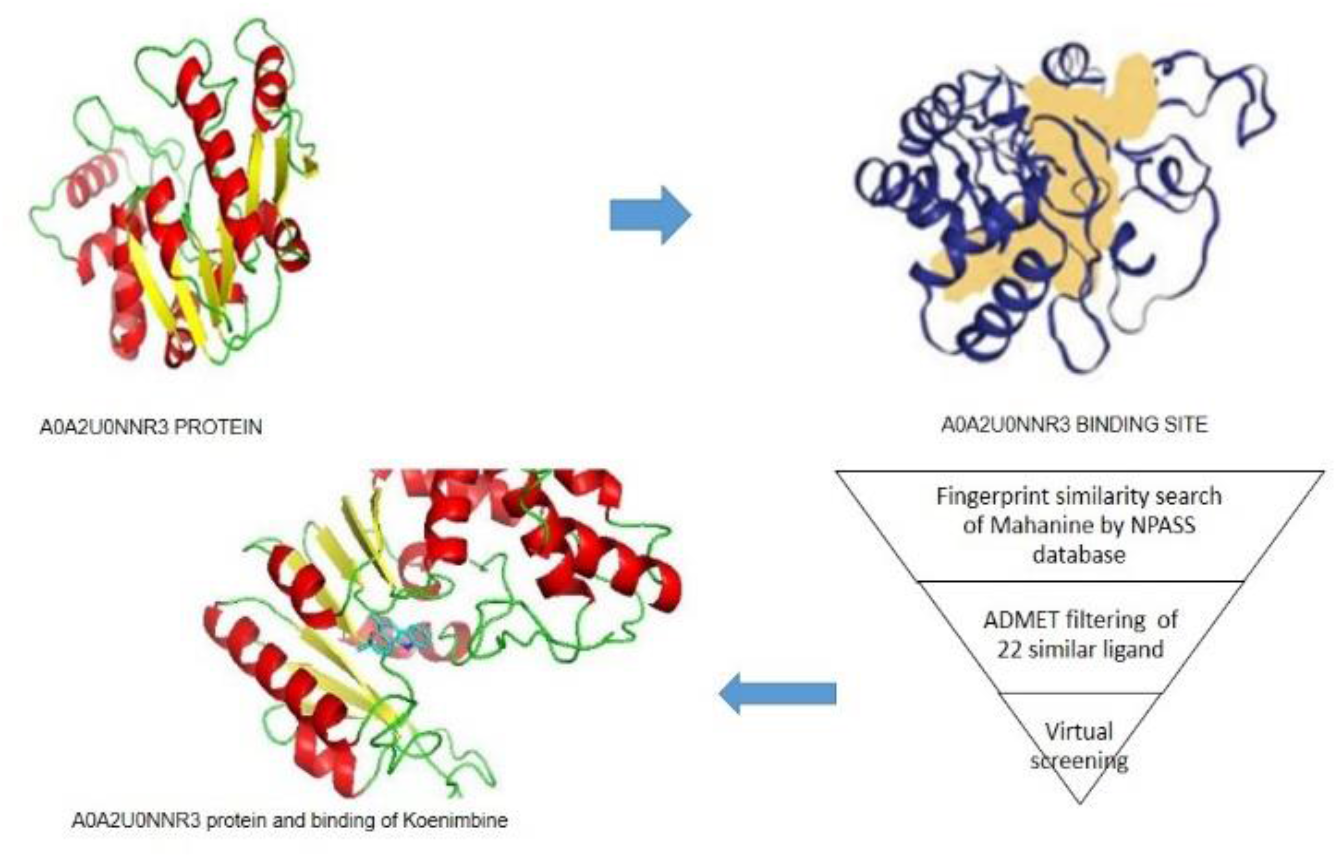
Figure shows the virtual screening method applied

### 2.1 Retrieval of bacterial proteome

All available strain *Klebsiella Pneumoniae* hypothetical protein sequence containing DNA adenine methyltransferase was obtained from UniProt (Universal Protein Database) by using keyword organisms; “*Klebsiella Pneumoniae”* and term; N6 DNA adenine methyltransferase and sequence containing unreviewed status are selected. The UniProt database is the largest protein database that is resourced with detailed annotation of protein (Bateman et al. 2017).

### 2.2 Identification of non-homology sequence analysis

All of the hypothetical protein sequences retrieved were first screened to the only select non-homologous sequence. The aim of this is to restrict and identify the hypothetical protein that is not similar to the human proteome. By performing BLASTp (Basic Local Alignment for protein) against the National Centre Biotechnology Institute database (NCBI). The threshold was set to an e-value of <0.0005(Anishetty, Pulimi, and Pennathur 2005).

### 2.3 Identification of essential genes

In order to identify a potential drug target, the non-homologous hypothetical protein that was first identified should consist of essential genes that are beneficial for the constituent cellular process of the cell. The essential genes are important to be identified as the major constituent to the survival of the pathogens. The hypoethetical proteins are screened by using BLAST against the Database of Essential Genes (DEG) and the threshold values were set to default parameters of E-value <0.0001 (Zhang, Ou, and Zhang 2004).

### 2.3 Broad-spectrum analysis

The hypothetical protein that contains all the required essential genes were then identified to be presented in a broad spectrum of the bacterial kingdom. Bacterial proteins are classified as broad-spectrum if the proteome is presented in more than 25 bacterial protein kingdoms. In order to identify these hypothetical proteins, the proteins of essential genes were screened by BLASTp with an E-value of 0.005.

### 2.4 Drugability analysis

DrugBank is extensively used for a new drug target in *silico* computational aided drug design (Hosen et al. 2015). The database consists of the data with more comprehensive information on the drug target and action of the drug. The drug data included in the database are approved by the Food and Drug Administration (FDA). In order to obtain the drugable protein, the hypothetical protein is subjected to BLASTp with an E-value of 0.001.

### 2.5 Non-homology analysis against gut microbiota

Gut microbiota plays an important role in human gastrointestinal. The homologous protein giving similarities to the human gut may interact and bind with the gut flora proteins leading to adverse pharmacokinetic side effects in the host. Hence, a homologous protein that is similar to the human gun is removed by using BLASTp with an e-value of 0.0001 (Altschul, Stephen F Gish, Warren Miller, Webb Myers 1990).

### 2.6 Subcellular localization

PSORTb 3.0 is an accurate predictor for bacterial protein subcellular localization(SCL). Gram-negative bacterial protein is composed of 5 major localization which includes cytoplasmic, inner membrane, periplasmic, outer membrane and extracellular (Yu et al. 2010). In this study, proteins that are located at the membrane channel and cytoplasmic are selected as they are more profound as a drug target (Barh et al. 2011). The subcellular localization identification is performed by using PsortB and the drug target located at cytoplasmic is selected.

### 2.7 Drug target property

Drugs are usually enzymes and they are involved in binding, signalling and communication. According to Bakheet (2009), good drug target are summarized to 8 major key properties namely, hydrophobicity >−142.4, length of amino > 550, signal motif present, no PEST motif, more than 2 *N*-glycosylated amino acids, not more than 1 O-glycosylated ser and mean pI of <7.2, presence of the transmembrane helix and membrane location is cytoplasmic. Therefore, all of these aspects are first analyzed to determine a good drug target.

To predict the length of amino acids, the ExPasy server is used to analyze the length of amino acids, hydrophobicity and pI. (Bendtsen et al. 2004). In order to identify the presence of transmembrane helix(THMM), the analysis is done via the TMHMM method (http://www.cbs.dtu.dk/~krogh/TMHMM/) (Krogh et al. 2001). PEST region is identified by via (http://emboss.cbr.nrc.ca/cgi-bin/emboss/epestfind). To analyze the O -glycosylation, The NetOglyc program via (http://www.cbs.dtu.dk/services/NetOGlyc/) is used while for N-glycosylation ser is accessed via (http://www.cbs.dtu.dk/services/NetNGlyc/) (L J Jensen et al. 2003).

### 2.8 Anti-target non-homology analysis

The anti-target non-homology analysis is performed to eliminate the anti-target receptor (Raman, Yeturu, and Chandra 2008). Anti-target proteins are eliminated with an E-value set to 0.005.

### 2.9 Drug data property

ChEMBL database provides the bioactivity, molecule, target and drug data from various data which includes the medicinal chemistry literature. The database provides an association for ligand and the specific biological target (Davies et al. 2015). The identified hypothetical protein was observed to show more matches from the ChEMBL as a good drug target.

### 2.10 Virulence factor analysis

Virulence Factor Database (VFDB) provides an extensive comprehension of the Virulence Factors that are characterized by 16 dominant bacterial pathogens (Chen et al. 2005). These virulence factors are one of the most crucial factors that cause bacterial pathogens to colonize the host and being detrimental to the host cell.

### 2.11 Protein-protein interaction

Protein functions are the main complex of the cellular phenotype and they are not independent. The networks of the interacting proteins elucidate the protein function. To obtain the protein-protein interaction (PPI) of *Klebsiella Pneumoniae*, STRING(Search Tool for the Retrieval of Interacting Genes/Protein) is used (Lars J Jensen et al. 2009). Neighboring protein with a high confidence score of greater than 0.7 is included.

### 2.12 Binding site prediction

Drugs bind to a specific site of the protein. The interaction between the binding site helps to understand the physicochemical interaction between the drugs and the protein and further interactions. The predictions are done by using DogSiteScorer, an automated algorithm for pocket and drug ability prediction. The pocket is predicted by mapping the protein to the grid and the Difference of Gaussian is used to filter and identify the pocket regions on the protein surface (Volkamer et al. 2012).

### 2.13 Metabolic Pathway Analysis

Comparative metabolic pathway analysis is performed to identify the unique interaction between the host and the identified protein by KEGG (Kyoto Encylopedia of Genes and Genomes). The output gives KO (KEGG Orthology) assignments that will generate the KEGG pathway(Kanehisa et al. 2010). Metabolic pathway analysis is essential to elucidate the predicted putative drugs.

### 2.14 Gene ontology

As the protein identified is uncharacterized protein, it is therefore essential to identify the specific biological role of the protein. Understanding the biological role of protein provides an insight into the protein specific function (Omeershffudin, and Suresh 2019). ThisThe biological role of the identified protein is accessed via Gene Ontology (GO). GO consortium is used for biology unification for all shared eukaryotes. The consortium constructs 3 ontologies which are the biological process, molecular function and the cellular component (Gene et al. 2011). The ontologies provide essential information to understand the biological role of the protein in specific organisms.

### 2.15 Homology modeling

Uncharacterized protein structure is not available in the Protein Data Bank (PDB) whilst the structural mechanism is vital to facilitate the understanding of the ligand interaction and also chanelling between the targeted protein (Kumar 2011a). The targeted 3D protein structure is constructed by using the fully automated server SWISS-MODEL. Firstly, homology modelling of targeted protein is first compared against a similar protein structure template (Kumar 2011b). The template for the identified protein was identified as the protein structure of bacteria *Escherichia Coli* containing the DNA adenine methyltransferase PDB ID: 4RTL(Horton et al. 2015).

The modeled protein structure is then verified to check the protein quality of the stereochemical by using PROCHECK (Laskowski, Molecular, and Thornton 1993), ProSA-web where errors in the 3D structure are recognized (Wiederstein and Sippl 2007), and ERRAT.

### 2.17 Ligand preparation

7 ligands are identified as DNA methyltransferase inhibitors (DNMTi) from the literature review (Saldívar-gonzález et al. 2018). The ligands are Mahanine (PubChem ID: 375151), Curcumin (PubChem ID: 969516), EGCG (PubChem ID: 65064), Nanaomycin A (PubChem ID: 40586), Parthenolide (PubChem ID: 7251185), Quercetin(PubChem ID: 5280343) and Trimethylaurintricarboxylic acid (PubChem ID: 263071). Ligands that can be developed for putative drug targets should not violate 5 Lipinski rules and therefore were first validify by the SWISS-ADME server. Ligands that results in any violation are not further included for analysis.

The 2D structure is obtained in SDF format and is retrieved from the PubChem database (https://pubchem.ncbi.nlm.nih.gov/.) The SDF file is converted to PDB via OpenBABEL and SMILES (http://cactus.nci.nih.gov/services/translate/) (Weininger 1988). The converted PDB structures are minimized to PDBQT with AutoDock Vina tools. Non-polar hydrogen is added and merges to the ligands and gaisteiger charges are computed. The torsion of the ligand is defined and saved as a PDBQT extension.

### 2.18 Molecular docking

Molecular docking is performed to identify ligands that bind with the lowest affinity score to develop potential putative drug targets by using AutoDock Vina. The default exhaustiveness is set to 1.0 Å. Identified protein is configured by first adding all hydrogens to the protein and merging the non-polar hydrogen. The gaisteiger charges are computed and the protein is saved as pdbqt file.

The grid box is set based on the predicted binding site with the configuration values of the center grid box of x,y,z. The size of the dimension grid box is set to 30.0 Å. The binding affinity score is observed.

### 2.19 Identification of novel inhibitor through molecular fingerprint search of the prioritized ligand

To identify novel inhibitor, fingerprint search is performed by NPASS(Natural Product Activity and Species Source databases) to search compound that is similar to the prioritized ligand based on docking of DNA methyltransferase inhibitors (Zeng et al. 2018). The fingerprint search is done using by inputing SMILES of a prioritized ligand with setting fingerprint type (pubchem-881 fp) with threshold >=0.90 search by structure function in NPASS database.

### 2.20 Virtual screening

Virtual screening is performed to dock against clusters of ligands by using AutoDock. The setting is set to default parameters of 1.0 Å with a dimension grid of 30.0 Å. The analysis is performed based on the binding energy score.

### 2.21 ADMET Test

Absorption, distribution, metabolism, excretion, and toxicity(ADMET) are the major aspects that are processed by the body as soon as the drug is administered into the body(Zhong 2017). These pharmacokinetic properties are accessed to indicate the site of action of the drugs. These aspects are analyzed by using the pkCSM database (Pires, Blundell, and Ascher 2015). PkCSM database provides optimization of these pharmacokinetics properties by using the graph-based signatures.

## 3.0 Result & Discussion

The aim of this study is to identify potential drug target among hypothetical proteins of *Klebsiella Pneumonia* containing DNA adenine methyltransferase and to identify novel DNMT inhibitors. All 32-hypothetical proteome of *Klebsiella Pneumoniae* containing DNA adenine Methyltransferase were retrieved from Uniprot to be analyzed as potential druggable protein. The druggable protein is first characterized by a subtractive genomics approach that characterizes protein of which suffices the following criteria; non-homologous to the human host, contains essential genes, presented in broad-spectrum bacterial kingdom, and non-homologous to the human gut microbiota.

The first consideration of the subtractive genomic approach is non-homology analysis. Homologous protein that is presented in the human host may react with the molecules as they may carry unwanted toxicity. To decrease the adverse side effects, non-homologous protein is selected as putative drug targets by subjecting the proteins sequence to BLAST with an E-value of 10- 3. Essential genes are known to be indispensable to maintain the cellular processes of the bacterial proteome to survive (Zhang, Ou, and Zhang 2004). To identify essential genes in the bacterial proteome, the sequences are subjected to BLASTp against the DEG database with an expected value of < 0.0001.

Threating multiple bacterial infections would require the target drug able protein to be common in the broad-spectrum bacterial kingdom(White and Kell 2004). Multiple target antibacterial is most preferred to be developed as drugs. To foretell whether these bacterial proteomes are broad-spectrum, all 32 protein is searched by performing BLAST with E-value of less than 0.0001 against NCBI bacterial pathogens database. This resulted in all 32 as non-homologous, contain essential genes and are presented in a broad spectrum of the bacterial kingdom.

### 3.1 Druggability analysis

Protein that is druggable is defined as being able to bind with drug molecules high strength of binding interaction (Wishart et al. 2008). These are known as high-affinity binding between the protein and ligand where it results in stronger attractions of intermolecular forces. One notable source containing comprehensive drug data is DrugBank. According to Hosen (2015), Drug Bank contains small molecule drugs, biotech drugs that are approved by FDA, nutraceuticals and experimental drugs entries. Targeted proteins are then BLASTp against the DrugBank database and out of 32 only 26 protein is characterized as being druggable.

### 3.2 Human gut microbiota analysis

The term gut microbiota is described as the large population of organisms that colonizes the intestinal tracts (Jandhyala et al. 2015). Gut microbiota is highly associated with human inflammatory diseases. The role of pathogens in human gut microbiota co-evolved by having a symbiotic relationship and promoting replication of pathogens (Joseph M. Pickard, Melody Y. Zeng, Roberta Caruso 2018). Homologous protein may lead to unintentional blockage of the proteins that are in the gut flora that could cause adverse effects (Shanmugham and Pan 2013). To prevent this, homologous protein is removed by BLASTp against the NCBI database of gut flora with a threshold of less than 0.005. Out of 26 protein, 19 protein is found to be non-homologous to the human gut.

### 3.3 Subcellular localization analysis

One important determinant of developing a putative drug target is the characterization of the subcellular localization as it exhibits the main function of the protein (Uddin et al. 2017). Localization of the protein determines the protein function (Chordia, Lakhawat, and Kumar 2017). Proteins that are located at cytoplasmic regions are seen to be more favorable as drug targets as they contain an abundant enzyme making it more feasible for a drug target. The cellular localization is predicted by using PsortB. 9 protein is characterized by locating at the cytoplasmic membrane while 10 protein is unknown.

### 3.3 Drug target property analysis

To further understand the drug property of this 9 protein, the protein is analyzed based on 8 major key properties summarized by Bakheet (2009). The 8 key properties of a good drug target are hydrophobicity >−142.4, length of amino > 550, a signal motif present, no PEST motif, more than 2 *N*-glycosylated amino acids, not more than 1 O-glycosylated ser and mean pI of <7.2, presence of the transmembrane helix and membrane location is cytoplasmic.

One of the driven forces of a good drug target is having high hydrophobicity. The balance of hydrophobicity of protein is significant to the folding of protein and aggregation(Zbilut et al. 2003). The higher the hydrophobicity of the protein, the better the resultant in the folding of the protein which indirectly aids in the functionality of the protein (Chordia, Lakhawat, and Kumar 2017). The stabilization of hydrophobicity of protein is also found to affect the interactions of the binding affinity between the protein-ligand (Snyder et al. 2011). The result shows that all protein has hydrophobicity of <−0.142.

Another observed key aspect of drug targets is the isoelectric point (pI) of the protein which is the overall charges of the amino acids. The pI value determines the pH of the protein and the protein solubility. Higher pI values indicate that the protein is basic and vice versa. The observed mean value of a good drug target is that the pI value should be <7.2. 3 proteins were identified as having a mean pI of <7.2: A0A3P4EC49, A0A3P4UG76, and A0A2U0NNR3. Table 1 shows the screening of the 3 proteins.

**Table 1:** Table 1 shows the results when performing the ChEMBL approach for identified targets and DogSiteScorer score pocket binding prediction of all 3 hypothetical proteins.

| Uniprot Id | ChEMBL -<br>targets identified | DoGSiteScorer |
| --- | --- | --- |
| **A0A3P4EC49** | 3 | 0.81 |
| **A0A3P4UG76** | 5 | 0.81 |
| **A0A2U0NNR3** | **10** | **0.82** |

The desirable amino acid length for drug target should be greater than 550 amino acids in total length. The greater the length of amino acids the greater the surface area of the protein interactions with drugs. Here, all the drug targets have less than 550 overall amino acids which may due to the type of the bacteria species. The signal peptide cleavage aids in the transportation of the protein of the endoplasmic reticulum across the membrane (Zimmermann et al. 2011). However, the result indicates that the presence of signalP is less significant due to the localization of the protein in the cytoplasmic.

PEST regions are regions of peptide that is rich with Proline(P), glutamic acid (E), Serine (S) and Threonine(T). Proteins that have one or more PEST regions are associated with having shorter intracellular half-lives as they are reported to cause protein degradation (Rogers, Wells, and Rechsteiner 1860). All of the protein sequences observed consist of at least 1 PEST region. The presence of transmembrane helix is amino acids that portray flanking regions however results show there well absent transmembrane helix in all observed protein.

Glycosylation is widely crucial and found abundant in polypeptide chain modification (Nothaft and Szymanski 2010). Bacterial protein possessed two glycosylation state which is the N-linked and O-linked glycosylation (Dell et al. 2010). According to Bakheet (2009), most bacteria protein drug targets either have greater than two N-glycosylated amino acids or one or no O-glycosylated ser. 4 proteins resulted in having one and no O linked glycosylated ser and 3 protein has more than 2 N linked glycosylated amino acids.

Based on the drug property 3 protein are selected to be further screened as they possessed more drug target characteristic which is: A0A3P4EC49, A0A3P4UG76, and A0A2U0NNR3.

### 3.4 Identification of putative drug target

It is necessary to prognosticate the probable drug candidate to fail. Therefore, the protein needs to first be identified as either an anti target or target protein. The anti target protein is a protein receptor that when it binds to a drug, it will cause adverse pharmacokinetics side effects. Here, all 3 proteins are not anti-target proteins.

Out of these 3 proteins, protein A0A2U0NNR3 is seen to have more desireable drug property when analyzed by using chEMBL with 10 matches. Figure 4 greatly summarizes the screening process to identify the potential drug target. Also, this protein shows the highest pocket binding score of 0.82 when analyzed by using DogSiteScorer. The protein then further studies for its metabolic pathway. Based on the metabolic pathway analysis in the KEGG server, the protein is found to carries a unique pathway of DNA mismatch repair. The gene ontology to describe the specific biological role of the protein indicates that the protein carries a gene that functions as a site-specific of DNA methyltransferase (adenine specific activity). The metabolic pathway and gene ontology are summarized in Table 2.

**Figure 4:**
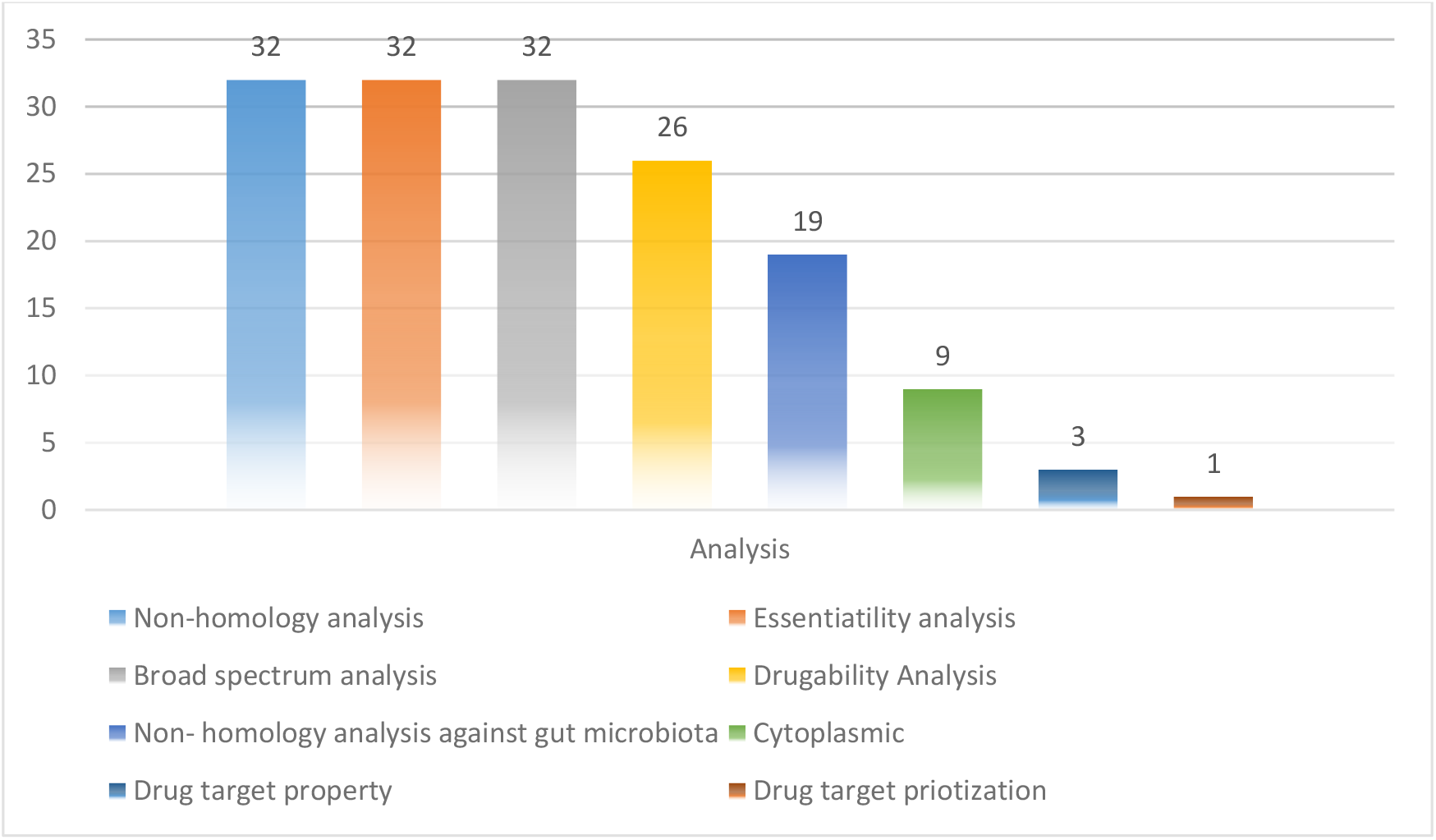
Analysis summarization of the subtractive genomic analysis of the 32-hypothetical protein of *Klebsiella Pneumoniae* containing DNA Adenine Methyltransferase.

**Table 2:**
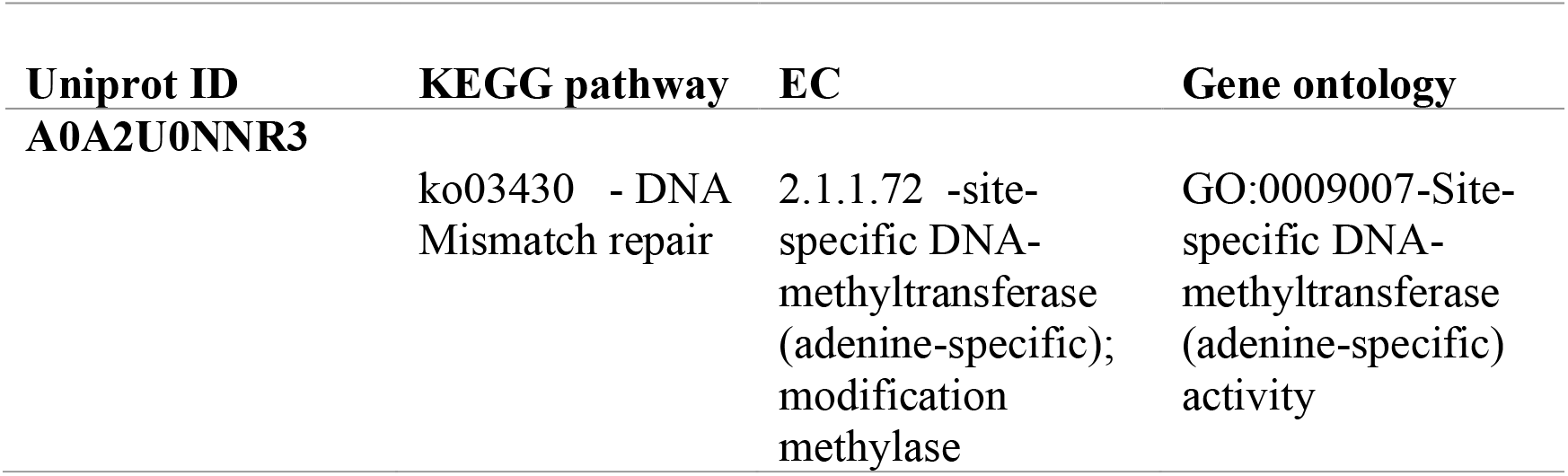
The KEGG pathway, EC number, and Gene ontology ID for protein A0A2U0NNR3

In order to understand the virulence mechanism, the protein is queried to Virulence Factor Databased. The finding resulted that the bacteria protein virulence factors are; VFG010749 (sdhB) Dot/Icm type IV secretion system effector, VFG001959 (hddC) capsular polysaccharide heptosyltransferas, VFG000166(pchE) dihydroaeruginoic acid synthetase PchE [Py], VFG000272 (ureE) urease accessory protein (ureE), metalloch and VFG002139 (cdsD) Type III secretion system inner membrane r. The virulence factor is summarized in Table 3.

**Table 3:** Virulence factor of the hypothetical protein of A0A2U0NNR3.

| UniProt ID | Virulence factors |
| --- | --- |
| A0A3P5S6Q8 | <u>VFG010749</u> (sdhB) Dot/Icm type IV secretion system effector |
|  | <u>VFG001959</u> (hddC) capsular polysaccharide heptosyltransferase |
|  | <u>VFG000166</u> (pchE) dihydroaeruginosic acid synthetase PchE [Py] |
|  | <u>VFG000272</u> (ureE) urease accessory protein (ureE), metalloch |
|  | <u>VFG002139</u> (cdsD) Type III secretion system inner membrane r |

### 3.4 Protein-protein interaction

Protein is not able to function alone, therefore the neighbouring protein is identified. Based on Figure 5, the interacting neighbouring protein helps to decipher the interactome mechanism in bacteria protein. Neighboring protein scored greater than 0.7 are considered as high confidence interacting protein which can be seen in Figure 6. The significant protein identified is tryptophanyl t-RNA synthetase, ribulose phosphate 3-epiramase, DNA mismatch repair endonuclease MutH, 3 dihydro quinate synthase, and DamX.

**Figure 5:**
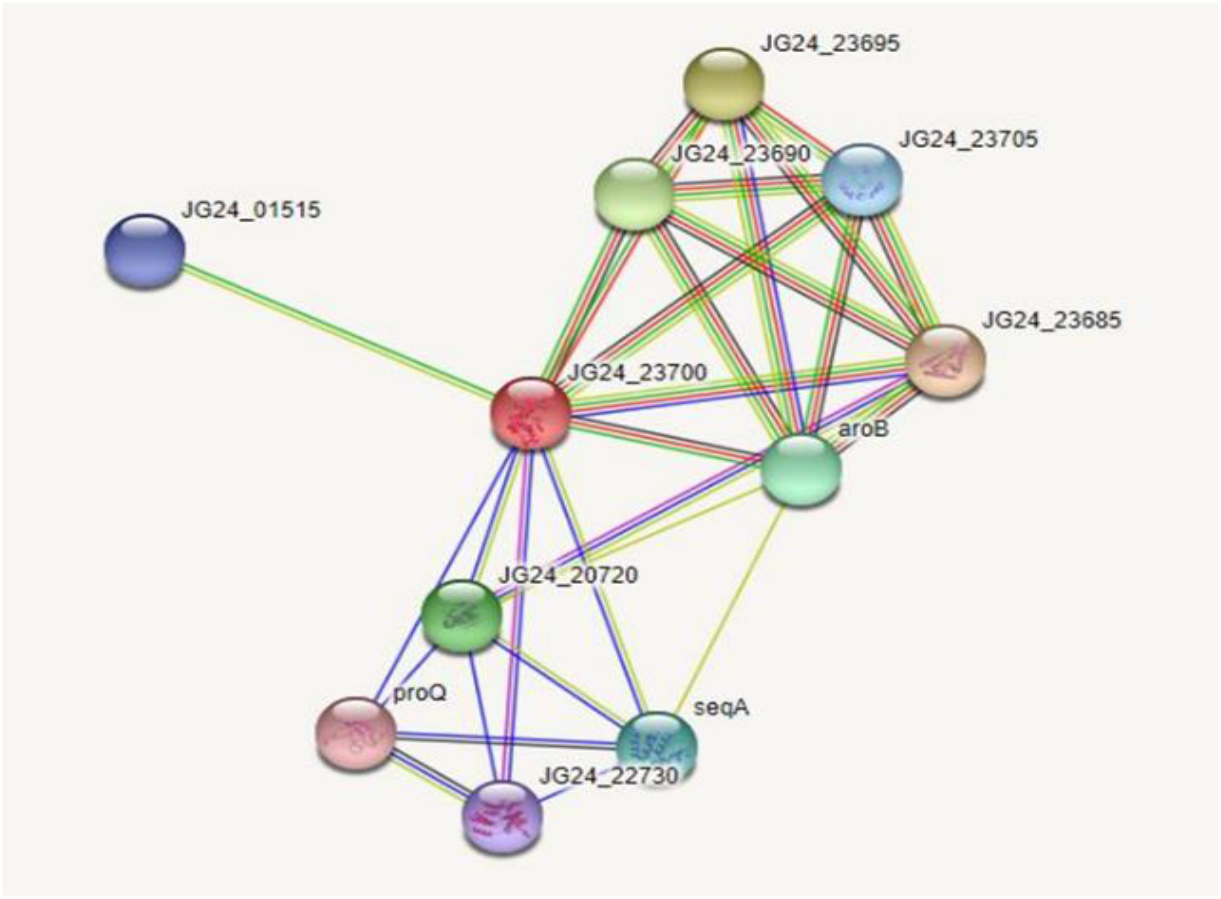
Shows the interacting protein between the main protein JG24_22730 of a specific site of DNA Adenine Methyltransferase and the functional partner

**Figure 6:**
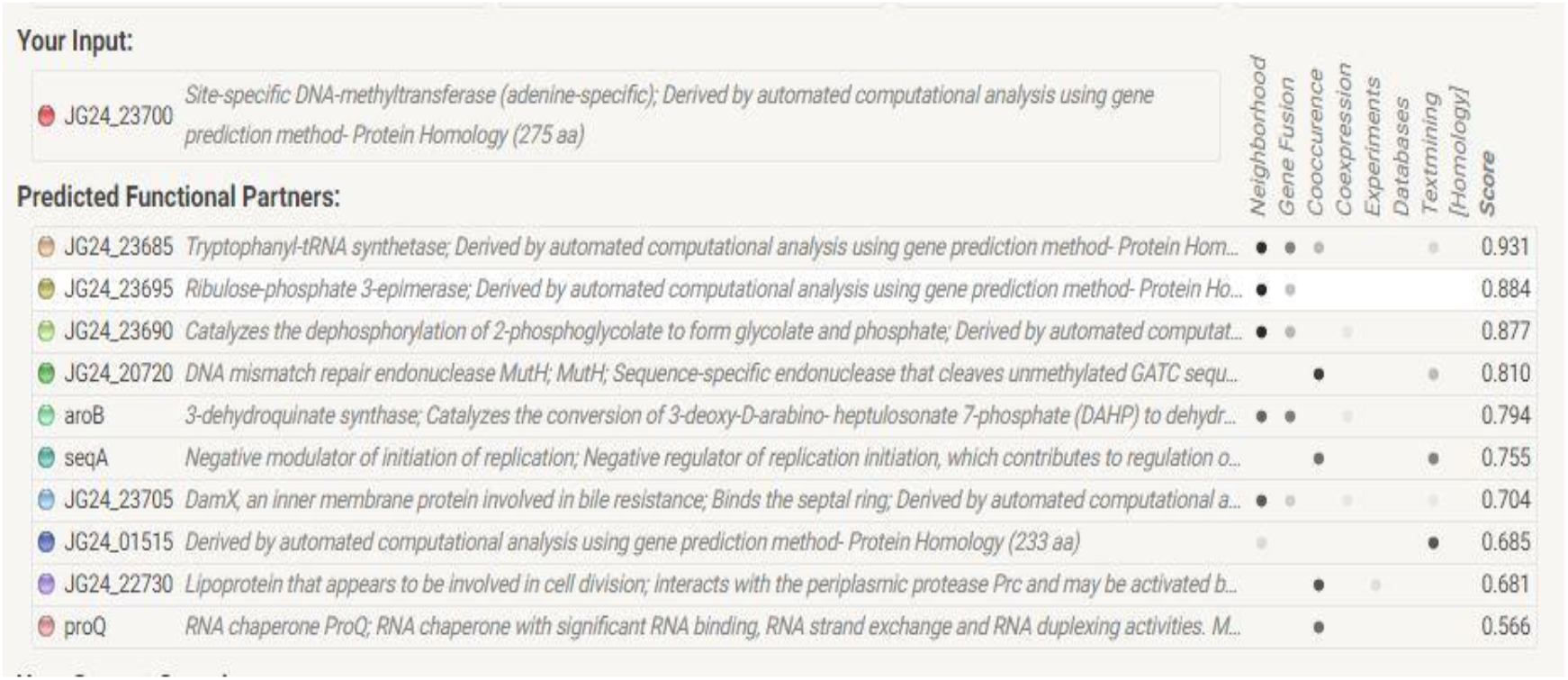
The detailed predicted functional partner of the protein interaction with the main protein of JG24_27300(Site-specific DNA-methyltransferase). The result demonstrates scores and of which the function is either; neighborhood, gene fusion, concurrence, and coexpression.

### 3.5 Molecular docking

Initially, the protein structure A0A2U0NNR3 is searched in Protein Data Bank. The search resulted that the protein structure is not available. The protein sequence is submitted to SWISS-MODEL for the construction of the protein structure. The modeled structure is imposed based on the retrieved template from the SWISS-MODEL template library of PDB ID: 4RTL. To ensure the structural quality of the model, the protein is first evaluated. The evaluation is done by using PROCHECK, ERRAT, and PRoSA. PROCHECK construct Ramachandran plot that gives 89.9% of residues in the favored regions, 9.4% regions in the additional allowed regions and 0.4% for both residues in generously and disallowed regions. According to Kumar (2015), good quality of protein structure should give an overall of residues in the favored and allowed regions of over 90%. Here, the model gives a total of 89.9% + 9.4% = 99.3%.

The Z-score value from ProSA is −7.49 that indicates that the model is by the X-ray crystallography method. ERRAT plot gives the overall factor quality score of 84.848 that gives an average quality score. The overall results can be referred to in Figure 8 and based on the results, the protein model can be inferred for further study. The ligands that are sourced from the literature review are first analyzed for any violations of Lipinski Rules. Only ECGC is excluded as it gives 2 violations of Lipinski Rules: N or O>10, NH or OH>5.

The inhibitors are docked and defined to which the predicted binding pocket by using DogSiteScorer. Figure 7 shows the area in which the active region is identified. The dimension grid is set to 30 Å. Based on the result tabulated in Table 4, Mahanine gives the highest binding affinity score of −10.8 kcal/mol followed by Nanaomycin A (−10.0 kcal/mol). Timethylaurintricarboxylic acid (−9.4 kcal/mol), Quercetin (−9.3 kcal/mol) and Curcumin give the lowest binding affinity score of −8.5 kcal/mol. The molecular docking result is tabulated in Table 4.

**Figure 7:**
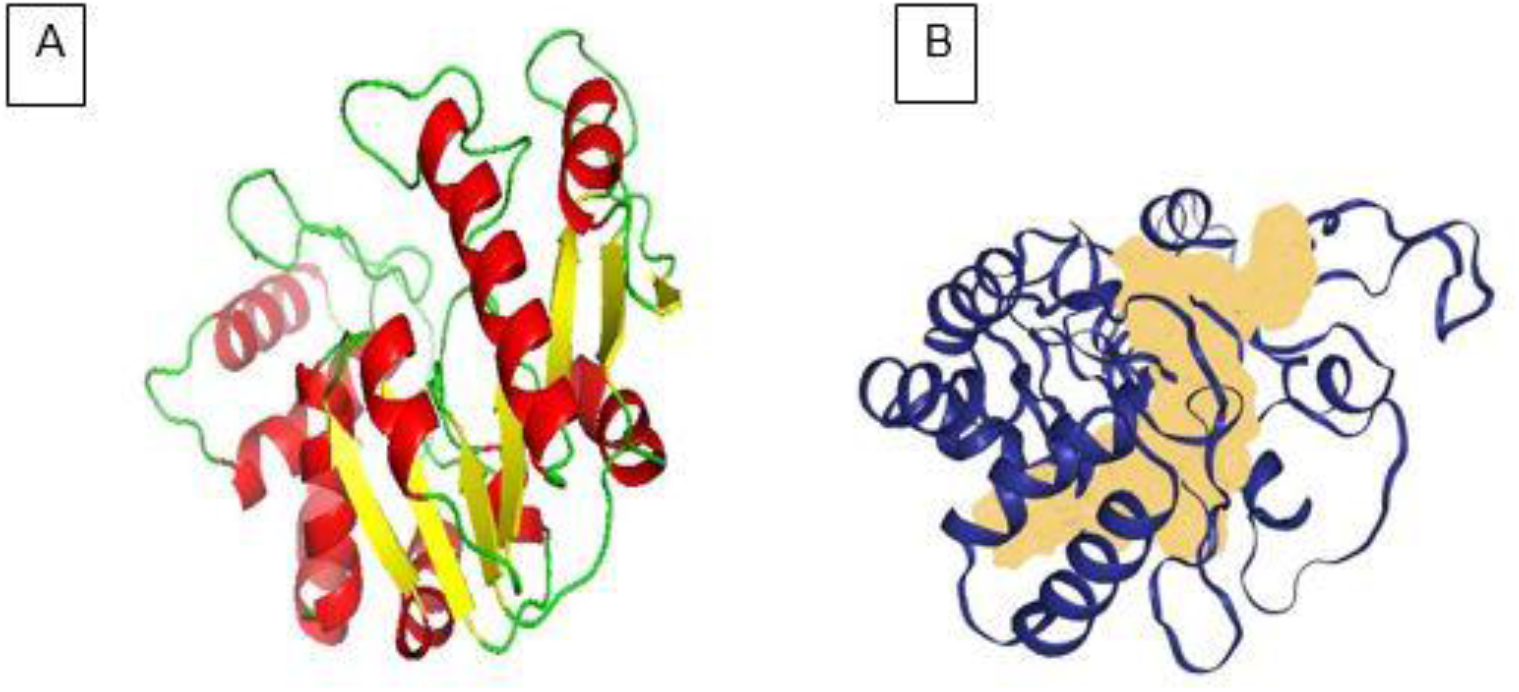
(A) Protein structure of A0A2U0NNR3 that is evaluated by using PyMOL. Red: helix, yellow: sheets and green: loops. (B) The active site that is demonstrated by the region colored peach is the binding site of the protein predicted by DogSiteScorer

**Figure 8:**
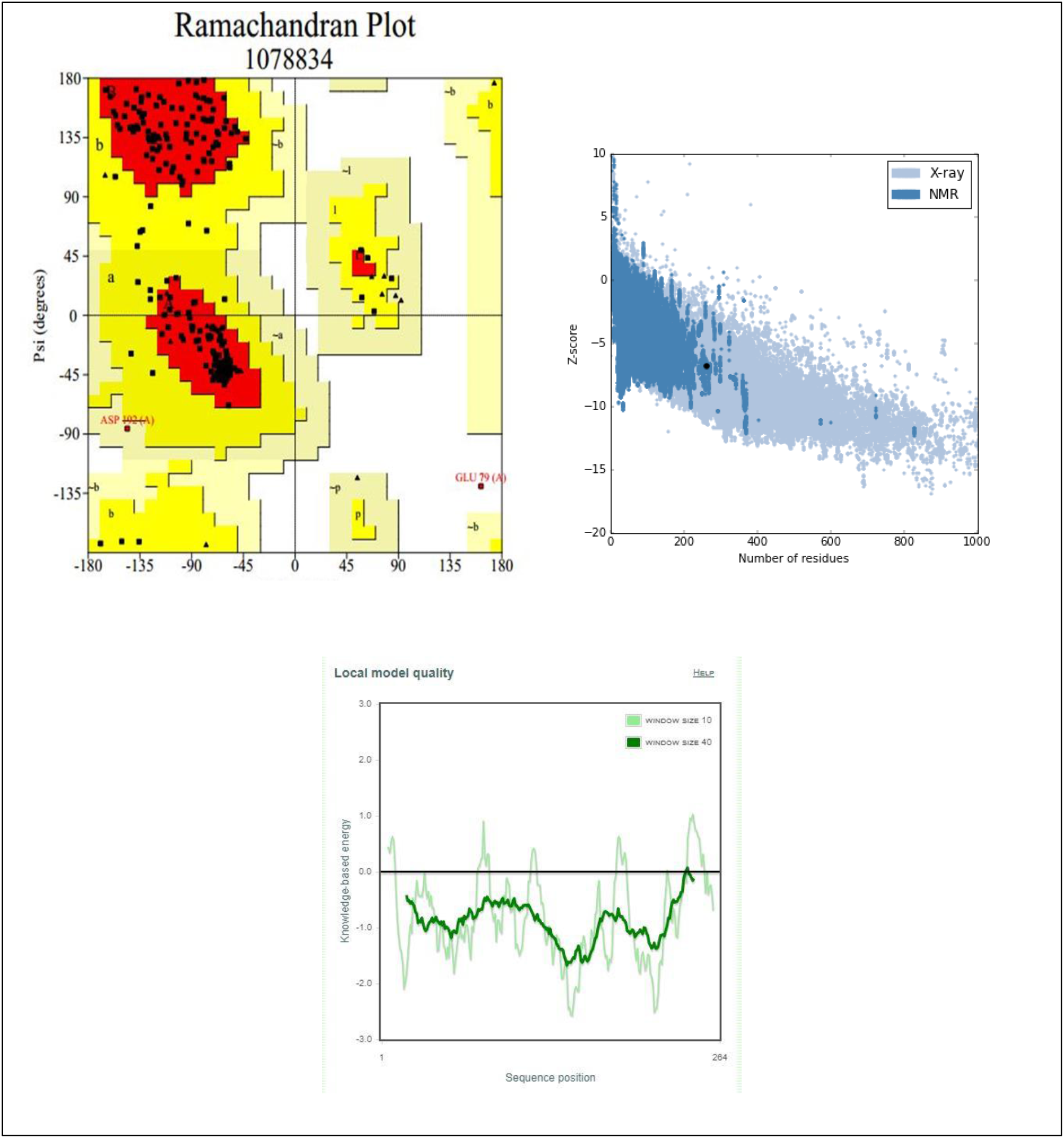
Structure quality assessment (A) Ramachandran plot of protein A0A2U0NNR3 by using SwissModel (B)Z-score of the residues (ProSA) (C) Local model quality based on the sequence position

**Table 4:** Binding affinity between the protein A0A2U0NNR3 against the 6 ligands

| Bioactive compound | Binding affinity(kcal/mol) |
| --- | --- |
| Curcumin | -8.5 |
| Mahanine | -10.8 |
| Nanaomycin A | -10.0 |
| Parthenolide | -7.6 |
| Quercetin | -9.3 |
| Trimethylaurintricarboxylic acid | -9.4 |

### 3.6 Virtual screening

Based on the binding affinity score, Mahanine is further screened to identify a compound with a similar composition as Mahanine. Mahanine is a carbazol alkaloid extracted from the plant species *Murraya Koenigii.* Mahanine is previously reported as one of the DNMT inhibitors. In order to identify new novel antibiotics derived from plant, fingerprint search is performed. Based on the calculated fingerprint similarities, 22 compounds are found to be similar to Mahanine. 3 compound name is not available. 2 compounds: Koenimbine and Grinibine are reported as one of the carbazole alkaloids of *Murraya Koenigii*. (Ramos et al. 2015).

The virtual screen result is tabulated in Figure 5.All the ligands are seen to no violate Lipinski Rules. The virtual screening is performed by using AutoDock. The virtual screening resulted that Koenimbine and Claurailas B have the highest binding affinity score of −5.97 kcal/mol. Koenimbine is also one of the carbazole alkaloids of the same plant species, can be explored as the novel antibiotics. Figure 9 shows the binding of Koenimbine and Table 5 shows the ligand-binding score and drug-likeness property.

**Figure 9:**
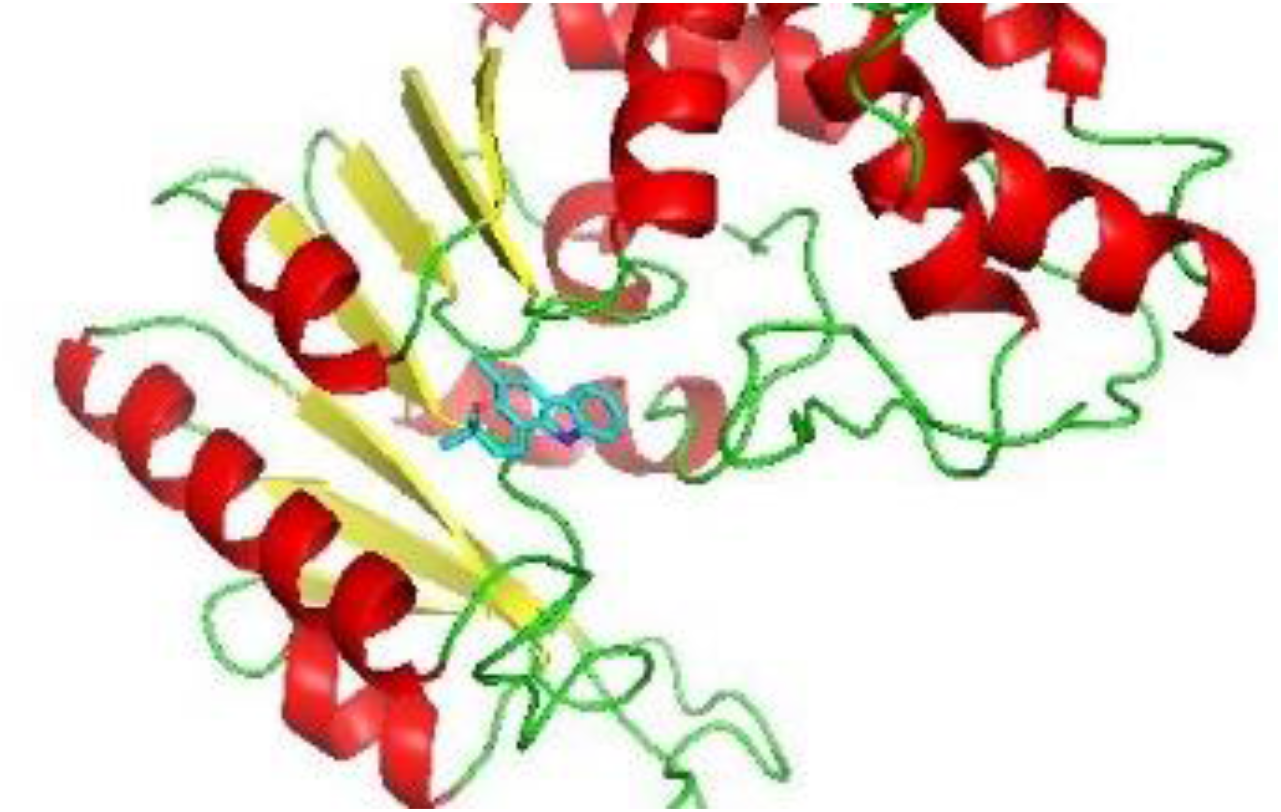
Binding of Koenimbine at the active site of protein A0A2U0NNR3

**Table 5:** Binding affinity score between the protein A0A2U0NNR3 against 22 ligands from similarity findings and drug-likeness property

| Natural product name | Product name | Pubchem ID | Binding energy Kcal/mol | Rotatable bonds | ADME TEST (Lipinski) |
| --- | --- | --- | --- | --- | --- |
| NPC162484 | Koenimbine | 97487 | -5.97 | 2 | Yes |
| NPC190007 | Mahanimbilol | 5353739 | -5.95 | 2 | Yes; 1 violations |
| NPC193777 | Glybomine B | 11208660 | -5.96 | 2 | Yes |
| NPC201697 | Clausenawalline C | 57409124 | -5.96 | 2 | Yes |
| NPC205934 | Mahanimbine | 167963 | -5.95 | 2 | Yes; 1 violation: MLOGP>4.15 |
| NPC209769 | Claurailas B | 51039826 | -5.97 | 2 | Yes |
| NPC212535 | 3-Methyl-9H-Carbazol-2-ol | 3459141 | -5.94 | 2 | Yes |
| NPC243834 | Sid506287 | 375146 | -5.95 | 2 | Yes |
| NPC267423 | n.a. | 375144 | -5.96 | 2 | Yes; 1 violation: MLOGP>4.15 |
| NPC291535 | Sid506290 | 375149 | -5.94 | 2 | Yes |
| NPC43787 | Grinimibine | 96943 | -5.92 | 2 | Yes |
| NPC470931 | n.a. | 71717470 | -5.89 | 2 | Yes; 1 violation: MLOGP>4.15 |
| NPC475085 | Sid506292 | 375151 | -5.91 | 2 | Yes |
| NPC475112 | Sid506289 | 375148 | -5.96 | 2 | Yes |
| NPC476044 | n.a. | 44583761 | -5.89 | 2 | Yes; 1 violation: MLOGP>4.15 |
| NPC476106 | 5-(3,5-Dimethyl-3,11-dihydropyrano[3,2-a]carbazol-3-yl)-2-methylpent-1-en-3-ol | 375155 | -5.96 | 2 | Yes |
| NPC477532 | 3-[(2E)-3,7-dimethylocta-2,6-dienyl]-8-[3-[(2E)- | 71725436 | -5.95 | 2 | No; 2 violations: MW>500, |
|  | 3,7-dimethylocta-2,6-dienyl]-9-methoxy-3,5-dimethyl-11H-pyrano[3,2-a]carbazol-10-yl]-3,5-dimethyl-11H-pyrano[3,2-a]carbazol-7-ol |  |  |  | MLOGP>4.15 |
| NPC48353 | Glycoborinine | 10446329 | -5.95 | 2 | Yes |
| NPC70259 | (+)-Mahanimbicine | 4072580 | -5.94 | 2 | Yes; 1 violation: MLOGP>4.15 |
| NPC70956 | Euchrestine B | 15060943 | -5.96 | 2 | Yes; 1 violation: MLOGP>4.15 |
| NPC72211 | (-)-O-Methylmahanine | 71716235 | -5.96 | 2 | Yes; 1 violation: MLOGP>4.15 |
| NPC94943 | Siamenol | 477436 | -5.95 | 2 | Yes |

### 3.8 ADMET test

The ADMET test was further evaluated for Koenimbine to understand the absorption, distribution, metabolism, excretion and toxicity of the compound. The analysis is analyzed and interpreted based on the guidelines of pkCSM (Pires, Blundell, and Ascher 2015). The compound absorption analysis resulted that the compound has high caco_2_ permeability. The water solubility is −4.618 log/mol/L. The compound absorbance in the human intestine is 93.479% which indicates a good absorption rate. It also results in being a P-glycoprotein II inhibitor which is significant in imposing pharmacokinetics effects.

In terms of distribution, koenimbine is seen to have slightly low distribution in the tissue plasma as the volume of distribution is 0.654 L/kg. Based on the toxicity measured, the *T.pyriformis* toxicity indicates that the compound is highly toxic against bacteria with 0.948 ug/L. The compound is also seen to be mutagenic against bacteria which indicates that it can cause detrimental to the bacteria. The result is summarized in Table 6.

**Table 6:** ADMET test for Koenimbine

| Absorption |  |  |  |  |  |  |
| --- | --- | --- | --- | --- | --- | --- |
| Water solubility<br>Numeric (log mol/L) | Caco2 permeability<br>Numeric (log Papp in 10<sup>-6</sup> cm/s) | Intestinal absorption (human)<br>(% Absorbed) | Skin permeability<br>Numeric (log Kp) | P-glycoprotein substrate | P-glycoprotein I inhibitor | P-glycoprotein II inhibitor |
| -4.618 | 1.411 | 93.479 | -2.742 | Yes | No | Yes |
| Distribution |  |  |  |  |  |  |
| VDss(human) |  | Fraction unbound (human) |  | BBB permeability (log BB) |  | CNS permeability (log PS) |
| 0.654 |  | 0.024 |  | 0.421 |  | -1.789 |
| Metabolism |  |  |  |  |  |  |
| CYP2D6 substrate | CYP3A4 substrate | CYP1A2 inhibitor | CYP2C19 inhibitor | CYP2C9 inhibitor | CYP2D6 inhibitor | CYP3A4 inhibitor |
| No | Yes | Yes | Yes | yes | No | No |
| Excretion |  |  |  |  |  |  |
| Total clearance (log ml/min/kg) |  |  | Renal OCT2 substrate |  |  |  |
| 0.477 |  |  | No |  |  |  |
| Toxicity |  |  |  |  |  |  |
| AMES toxicity |  | Max tolerated dose (human)<br>(log mg/kg/day) | Hepatotoxicity | Skin sensitization |  | Minnow toxicity (log mM) |
| Yes |  | -0.312 | Yes | No |  | -0.436 |

## 4.0 CONCLUSION

We identified and prioritized one DAM drug target from 32 proteins of *Klebsiella pneumonia* based on subtractive genomics, based on drug property, pocket analysis, pathway analysis, and structure analysis. Our proposed new computational pipeline approach helps to find out the drug target in a rapid way. We have also discovered that Koenimbine a natural bioactive chemical compound of *Murraya Koenigii* can potentially be a novel antibiotic We have also discovered that Mahanine can be used as a potential novel antibiotic to inhibit DAM containing bacterial pathogens.

## Authors Contribution

Umairah Natasya Mohd Omeershffudin: Conceptualization, Investigation, Methodology, Formal analysis, Data curation, Writing - original draft. Suresh Kumar: Conceptualization, Data curation, Validation, Visualization, Project administration, Resources, Supervision, Roles/Writing - review & editing

## ACKNOWLEDGMENT

The authors would like to thank the Faculty of Health Science, Management and Science University for supporting facilities to carry out the research work.

